# Concurrent tau pathologies in frontotemporal lobar degeneration with TDP-43 pathology

**DOI:** 10.1101/2021.09.23.461523

**Authors:** Shunsuke Koga, Xiaolai Zhou, Aya Murakami, Cristhoper Fernandez De Castro, Matthew C. Baker, Rosa Rademakers, Dennis W. Dickson

**Affiliations:** Department of Neuroscience, Mayo Clinic, Jacksonville, Florida, USA; State Key Laboratory of Ophthalmology, Zhongshan Ophthalmic Center, Sun Yat-Sen University, Guangzhou, Guangdong, China; Applied and Translational Neurogenomics, VIB Center for Molecular Neurology, VIB, Antwerp, Belgium; Department of Biomedical Sciences, University of Antwerp, Antwerp, Belgium

## Abstract

**Aims:** Accumulating evidence suggests that patients with frontotemporal lobar degeneration (FTLD) can have pathologic accumulation of multiple proteins, including tau and TDP-43. This study aimed to determine the frequency and characteristics of concurrent tau pathology in FTLD with TDP-43 pathology (FTLD-TDP).

**Methods:** The study included 146 autopsy-confirmed cases of FTLD-TDP and 55 cases of FTLD-TDP with motor neuron disease (FTLD-MND). Sections from the basal forebrain were screened for tau pathology with phospho-tau immunohistochemistry. For cases with tau pathology on the screening section, additional brain sections were studied to establish a diagnosis. Genetic analysis of *C9ORF72*, *GRN*, and *MAPT* was performed on select cases.

**Results:** Among 201 cases, we found 72 cases (36%) with primary age-related tauopathy (PART), 85 (42%) with aging-related tau astrogliopathy (ARTAG), 45 (22%) with argyrophilic grain disease (AGD), and 2 cases (1%) with corticobasal degeneration (CBD). Patients with ARTAG or AGD were significantly older than those without these comorbidities. One of the patients with FTLD-TDP and CBD had *C9ORF72* mutation and relatively mild tau pathology, consistent with incidental CBD.

**Conclusion:** The coexistence of TDP-43 and tau pathologies was relatively common, particularly PART and ARTAG. Although rare, individual patients with FTLD can have multiple concurrent proteinopathies. The absence of TDP-43-positive astrocytic plaques may suggest that CBD and FTLD-TDP were independent disease processes in the two patients with both tau and TDP-43 pathologies. It remains to be determined if mixed cases represent a unique disease process or two concurrent disease processes in an individual.

## Introduction

Frontotemporal lobar degeneration (FTLD) is pathologically and clinically heterogeneous and classified by the predominant protein that accumulates within neuronal and glial lesions. Most cases of FTLD have either transactive response DNA-binding protein of 43 kDa (TDP-43; FTLD-TDP, 50%) or tau inclusions (FTLD-tau, 45%), with a small number having inclusions of fused in sarcoma (FUS; FTLD-FUS, <5%) [1]. Patients with FTLD-TDP have heterogenous clinical presentations, including behavioral variant frontotemporal dementia, primary progressive aphasia, and corticobasal syndrome. Familial FTLD-TDP is most commonly caused by mutations in *C9ORF72* or *progranulin* (*GRN*) [2]. Some patients with FTLD also develop motor neuron disease (MND; FTLD-MND). Accumulation of TDP-43 aggregates in the central nervous system is a common pathologic feature of both MND and FTLD; thus, both are considered part of a spectrum of TDP-43 proteinopathies.

FTLD-tau includes progressive supranuclear palsy (PSP), corticobasal degeneration (CBD), and Pick’s disease [3]. Intracellular aggregates of phosphorylated tau protein in neurons and glia associated with neurodegeneration are pathologic hallmarks of FTLD-tau [3]. Increased age is a common risk factor for neurodegenerative disorders. Age-related changes, such as cellular senescence, mitochondrial dysfunction, and epigenetic alterations, have been reported in neurodegenerative processes [4–6]; thus, elderly individuals may often develop more than one neurodegenerative disease, with the accumulation of multiple types of pathological protein aggregates [7–10]. Indeed, concurrent TDP-43 pathology has been reported in a range of tauopathies, including Alzheimer’s disease [10, 11], PSP [12–14], and CBD [14–17]. Our previous study found that 45% of CBD patients had TDP-43 pathology [17]. Some cases had extensive TDP-43 pathology in the neocortex, which can be considered a mixed FTLD-tau and FTLD-TDP. More recently, Kim and colleagues reported nine cases of mixed FTLD-TDP and FTLD-tau, in which three unclassifiable FTLD-tau and two PSP cases had a primary diagnosis of FTLD-TDP, while FTLD-TDP was found in four cases with CBD [18]. These studies suggest that FTLD can be caused by coexisting TDP-43 and tau pathologies; however, the frequency and characteristics of tau and TDP-43 copathology have not been investigated.

In the present study, we aimed to determine the frequency and characteristics of tau pathology in a series of cases of FTLD-TDP and FTLD-MND that were initially considered “primary” TDP-43 proteinopathies as their original neuropathologic diagnosis. To do this, we screened tau pathology in 146 patients with FTLD-TDP and 55 patients with FTLD-MND. All cases were from the Mayo Clinic brain bank for neurodegenerative disorders.

## Methods

### Case selection

This study included 201 cases with FTLD-TDP with (N = 55) or without MND (N=146) in the period of 1998 to 2020. All brain autopsies were performed after consent of the legal next-of-kin or individual with power-of-attorney to grant permission. Studies of autopsy samples are considered exempt from human subject research by Mayo Clinic Institutional Review Board.

### General neuropathologic assessment

Formalin-fixed brains underwent systematic and standardized sampling with neuropathologic evaluation by a single, experienced neuropathologist (D.W.D.). Regions sampled in all cases included six regions of the neocortex, two levels of hippocampus, a basal forebrain section (including amygdala, lentiform nucleus and hypothalamus), corpus striatum at level of the nucleus accumbens, thalamus at the level of the subthalamic nucleus, midbrain, pons, medulla, and two sections of cerebellum, one including the deep nuclei. Paraffin-embedded 5-μm thick sections mounted on glass slides were stained with hematoxylin and eosin (H&E) and thioflavin S (Sigma-Aldrich, St. Louis, MO). Braak neurofibrillary tangle stage (NFT), Thal amyloid phase, and severity of cerebral amyloid angiopathy were assigned using thioflavin S fluorescent microscopy according to previously described methods [19–23].

Immunohistochemistry for phospho-TDP-43 (pS409/410, mouse monoclonal, 1:5000, Cosmo Bio, Tokyo, Japan) was performed on sections of cortex, hippocampus, basal forebrain, midbrain, medulla, and cervical spinal cord to establish a neuropathological diagnosis of FTLD-TDP or FTLD-MND. The neuropathologic diagnosis of FTLD-MND required motor neuron loss with Bunina bodies and a variable degree of corticospinal tract degeneration, demonstrated with myelin stains (Luxol fast blue-periodic-Schiff) and immunohistochemistry for activated microglia (IBA-1, rabbit IG, 1:3000, Wako Chemicals, USA [24]). Hippocampal sclerosis was diagnosed when neuronal loss and gliosis were selective in the CA1 sector and/or subiculum of the hippocampus in the absence of other pathologic findings that could account for neuronal loss in this region.

### Screening of tau pathologies

We immunostained 5-μm-thick sections of the basal forebrain section using a phosphorylated-tau antibody (phospho-tau Ser202, CP13; mouse monoclonal; 1:1000; a gift from the late Dr. Peter Davies; Feinstein Institute for Medical Research). Following deparaffinization in xylene and reagent alcohol, antigen retrieval was performed by steaming slides in distilled water for 30 minutes. Immunostaining was performed with an IHC Autostainer 480S (Thermo Fisher Scientific Inc., Waltham, MA) and DAKO EnVision™ + reagents (Dako, Carpinteria, CA). Immunostained slides were counterstained with hematoxylin and coverslipped. Tau-immunostained slides of all 201 cases were evaluated by three investigators (SK, XZ, DWD) blinded to clinical and pathological information. For cases suspected of having FTLD-tau based on the basal forebrain screening section, additional sections from motor cortex, cingulate gyrus, superior frontal gyrus, thalamus/subthalamic nucleus, midbrain, pons, and cerebellum were also processed for tau immunohistochemistry.

A diagnosis of CBD was made based on presence of astrocytic plaques and numerous tau-positive threads in the gray and white matter in cortical and subcortical regions [25]. A diagnosis of argyrophilic grain disease (AGD) required tau-positive (argyrophilic) grains in medial temporal lobe structures (i.e., amygdala and hippocampus), accompanied by pretangles, coiled bodies, balloon neurons, and granular/fuzzy astrocytes or bush-like astrocytes [26]. Silver stains (Gallyas) were used to confirm the diagnosis of AGD. For cases with AGD in the amygdala, additional sections from hippocampus, entorhinal cortex, inferior temporal gyrus, and cingulate gyrus were stained with tau immunohistochemistry and assessed to assign an AGD stage according to Saito et al. [27]. The diagnosis of aging-related tau astrogliopathy (ARTAG) was associated with variable thorn-shaped astrocytes or granular/fuzzy astrocytes in subependymal, subpial, perivascular, gray matter, and white matter [28].

### Clinical assessment

Clinical information was abstracted by two investigators (SK and AM) from the available medical records and brain bank questionnaires filled out by a close family member. The information included the age at symptom onset, disease duration, age at death, clinical diagnosis, clinical symptoms, neurological signs, and family history of dementia or parkinsonism.

### Genetic Analysis

We performed genetic analyses in two patients with FTLD-TDP and CBD. For genotyping, genomic DNA was extracted from frozen cerebellum tissue using standard procedures. *MAPT* H1/H2 haplotype (SNP rs1052553 A/G, A = H1, G = H2) was assessed with TaqMan SNP genotyping assays (Applied Biosystems, Foster City, CA). Genotype calls were obtained with QuantStudio™ Real-Time PCR Software (Applied Biosystems). *MAPT* sequencing was performed in exons 7, 9, 10, 11, 12, and 13, as well as known pathogenic intronic mutations located at 50 bp on either side of each exon (e.g., IVS10+16 C>T). *GRN* sequencing and screening for *C9ORF72* hexanucleotide repeat expansion were performed as previously described [29, 30].

#### Statistical Analysis

All statistical analyses were performed using R 3.4.3. Fisher’s exact test was performed for group comparisons of categorical data, as appropriate. Mann-Whitney rank-sum test, analysis of variance (ANOVA) on ranks, followed by Steel-Dwass post hoc test, or one-way ANOVA, followed by post hoc Tukey test, was used for analyses of continuous variables as appropriate. P-values <0.05 were considered statistically significant.

## Results

### Summary of the cohort

The study set included 146 patients (84 men and 62 women) of FTLD-TDP and 55 patients (30 men and 25 women) of FTLD-MND (**Table 1**). The average age at death was significantly older in FTLD-TDP than in FTLD-MND (74 ± 10 vs. 67 ± 9 years; p<0.001). FTLD-TDP also had a significantly longer disease duration than FTLD-MND (9 ± 5 vs. 4 ± 2 years; p<0.001). The average formalin-fixed brain weight was significantly lower in FTLD-TDP than in FTLD-MND (970 ± 160 g vs. 1120 ± 170 g grams; p<0.001). Alzheimer-type pathologies, measured by the Braak NFT stage and Thal amyloid phase, were not significantly different between the two groups.

**Table 1:** Case series and frequency of tau pathology among disease groups.

|  | **Overall**<br>**N = 201** | **FTLD-TDP**<br>**N = 146** | **FTLD-MND**<br>**N = 55** | **P value** |
| --- | --- | --- | --- | --- |
| Male% (N) | 57% (195) | 58% (84) | 55% (30) | 0.449 |
| Age, year | 72 ± 10 | 74 ± 10 | 67 ± 9 | 6.5 x 10<sup>-7</sup> |
| Disease duration, year | 7 ± 5 | 9 ± 5 | 4 ± 2 | 1.9 x 10<sup>-10</sup> |
| Brain weight, gram | 1010 ± 180 | 970 ± 160 | 1120 ± 170 | 1.3 x 10<sup>-8</sup> |
| Braak NFT Stage | II (I, II) | II (I, III) | II (I, II) | 0.707 |
| Thal phase | 0 (0, 2) | 1 (0, 2) | 0 (0, 2) | 0.400 |
| CAA | 31% (64) | 34% (49) | 25% (14) | 0.308 |
| PART | 36% (72) | 34% (50) | 40% (22) | 0.540 |
| ARTAG | 42% (85) | 44% (64) | 39% (21) | 0.629 |
| AGD | 22% (45) | 23% (33) | 22% (12) | 1.000 |
| CBD | 1% (2) | 1% (2) | 0% (0) | 1.000 |

Abbreviations: AD, Alzheimer's disease; AGD, argyrophilic grain disease; ALS, amyotrophic lateral sclerosis; ARTAG, aging-related tau astrogliopathy; CAA, cerebral amyloid angiopathy; PART, primary age-related tauopathy; FTLD, Frontotemporal lobar degeneration; MND, motor neuron disease.

### Alzheimer’s-type pathology assessed by thioflavin S microscopy

Thioflavin S microscopy was used to assign Braak NFT stage: Braak stage 0 in 46 (23%), stage I in 40 (20%), stage II in 65 (32%), stage III in 28 (14%), stage IV in 14 (7%), stage V in 4 (2%), and stage VI in 4 patients (2%). Thal amyloid phase was also assigned using Thioflavin S microscopy: Thal phase 0 in 106 (53%), phase 1 in 41 (20%), phase 2 in 23 (11%), phase 3 in 12 (6%), phase 4 in 7 (3%), and phase 5 in 12 (6%). Cerebral amyloid angiopathy was detected in 64 patients (61%). In this autopsy cohort, 72 patients (36%) had Braak stages I - IV and Thal amyloid phase 0, consistent with a diagnosis of primary age-related tauopathy (PART). The age at death was not significantly different between patients with and without PART (71 ± 9 vs. 73 ± 10 years; 0.194).

### Frequency of tau pathologies in TDP-43 proteinopathies

The second most common concurrent tau pathology was ARTAG, which was detected in 85 cases (42%) (**Table 1**). On tau immunohistochemistry, thorn-shaped astrocytes were detected in subpial (**Figure 1A**) and perivascular spaces (**Figure 1B**) in the mediobasal forebrain, and variably in the amygdala (**Figure 1C**). Thorn-shaped astrocytes were also observed in the peri-amygdaloid white matter (**Figure 1D**). The age at death was significantly older in cases with ARTAG than in those without ARTAG (76 ± 9 vs. 69 ± 10 years; p = 5.7 x 10^−7^), regardless of the disease group (i.e., FTLD-TDP, FTLD-MND; **Figure 1E**). A multivariable logistic regression model adjusting for age, sex, Braak NFT stage, and Thal amyloid phase revealed that older age (OR 1.08; CI 1.04-1.12; p = 3.4 x 10^−5^), higher Braak NFT stage (OR 2.39; CI 1.24-4.61; p = 0.010), and male sex (OR 1.33; CI 1.05-1.69; p = 0.018) were independent risk factors for ARTAG.

**Figure 1:**
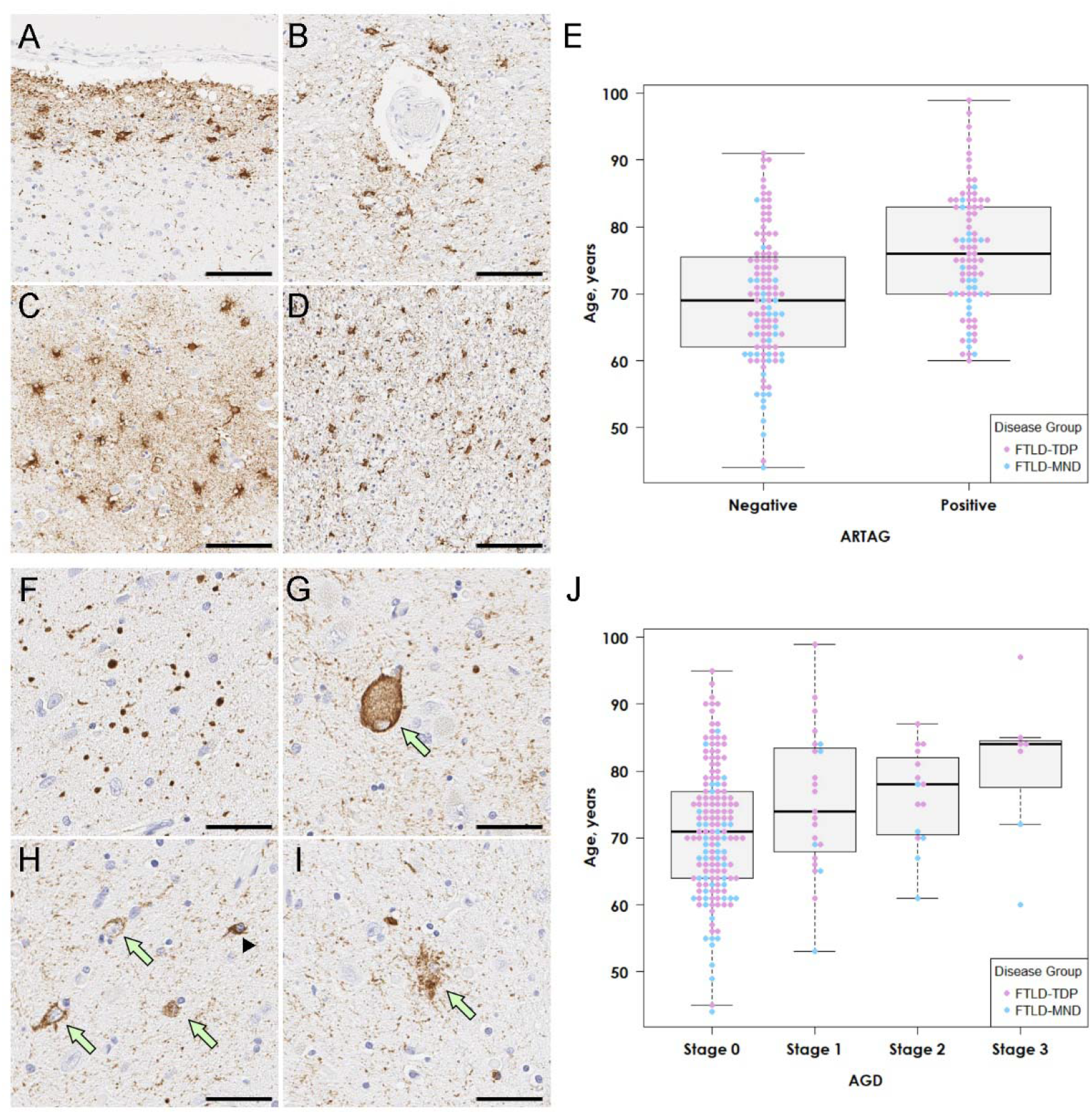
Representative images of ARTAG and AGD. (A-D, F-I) Immunohistochemistry for CP13. (A-D) Numerous thorn-shaped astrocytes are observed in the subpial mediobasal region (A) and perivascular regions (B), and the gray matter (C) in the amygdala. Thorn-shaped astrocytes are also present in the peri-amygdaloid white matter (D). (E) Patients with ARTAG are significantly older than those without ARTAG across the three diseases. (F-I) The amygdala has numerous argyrophilic grains (F), along with balloon neuron (G), pretangles (H, arrows), coiled bodies (H, arrowhead), and granular-fuzzy astrocytes (I), consistent with AGD. (J) Patients with AGD (stages 1, 2, and 3) are significantly older than those without AGD (stage 0) across the three diseases. Scale bars: 50 μm in A-D and F-I.

AGD was detected in 45 cases (22%). Argyrophilic grains in the amygdala were accompanied by pretangles, coiled bodies, balloon neurons, and GFA (**Figure 1F–I**). Screening of additional regions revealed that 23 cases had AGD restricted to the amygdala (stage 1), 15 also had AGD in the entorhinal cortex or subiculum (stage 2), and in 7 cases had pathology in the cingulate gyrus (stage 3). As with ARTAG, the age at death was significantly older in cases with AGD compared to cases without AGD (77 ± 10 vs. 71 ± 10 years; p = 9.1 x 10^−4^). As shown in **Figure 1J**, the median age at death was highest in stage 3, followed by stages 2 and 1. A multivariable logistic regression model adjusting for age, sex, Braak NFT stage, and Thal amyloid phase revealed that older age (OR 1.05; CI 1.01-1.09; p =0.017), and higher Braak NFT stage (OR 1.33; CI 1.03-1.71; p = 0.026) were independent risk factors for AGD.

We found two patients with tau pathology consistent with CBD (**Table 2**). Immunohistochemistry for tau revealed astrocytic plaques in the superior frontal gyrus and premotor cortex, tau-positive threads and coiled bodies in the adjacent white matter, and pretangles and threads in the subthalamic nucleus, pontine base, inferior olivary nucleus, and cerebellar dentate nucleus in both patients (**Table 3**). One patient (Case 2) also had AGD. The other patient who had a family history of dementia (Case 1) carried *C9ORF72* hexanucleotide repeat expansion. Mutations in *GRN* or *MAPT* were not detected in either patient. Clinical presentations of these patients were Alzheimer’s type dementia (Case 1) and behavioral variant FTD (Case 2). To adjudicate which pathology was most likely to contribute to their clinical presentations, we describe detailed pathologic findings and clinical features of these two cases. The detailed clinical history is provided in **Supplementary Data**.

**Table 2:** Clinicopathologic features of mixed FTLD-TDP and CBD cases.

|  | Case 1 | Case 2 |
| --- | --- | --- |
| Clinical diagnosis | Familial AD | FTD |
| Sex | Male | Male |
| Age at death, year | 89 | 84 |
| Age at onset, year | 82 | 78 |
| First symptoms | Memory loss | Memory loss,<br>behavioral changes |
| Family history of dementia or<br>parkinsonism | + | - |
| Motor weakness | - | - |
| Parkinsonism | + | - |
| Cognitive impairment | + | + |
| Brain weight, gram | 920 | 1080 |
| Braak NFT stage | III | I |
| Thal amyloid phase | 3 | 1 |
| Thal CAA stage | 2 | 0 |
| TDP-43 pathology | FTLD-TDP A;<br>HpScl | FTLD-TDP A;<br>HpScl |
| Tau pathology | CBD | CBD; stage 3 AGD |
| *MAPT* haplotype | H1/H1 | H1/H1 |
| *MAPT* mutations | Negative | Negative |
| *C9ORF72* mutations | Positive | Negative |
| *GRN* mutations | Negative | Negative |

Abbreviations: AD, Alzheimer's disease; AGD, argyrophilic grain disease; CAA, cerebral amyloid angiopathy; CBD, corticobasal degeneration; FTD, frontotemporal dementia; FTLD-TDP, frontotemporal lobar degeneration with TDP-43; HpScl, hippocampal sclerosis.

**Table 3:**
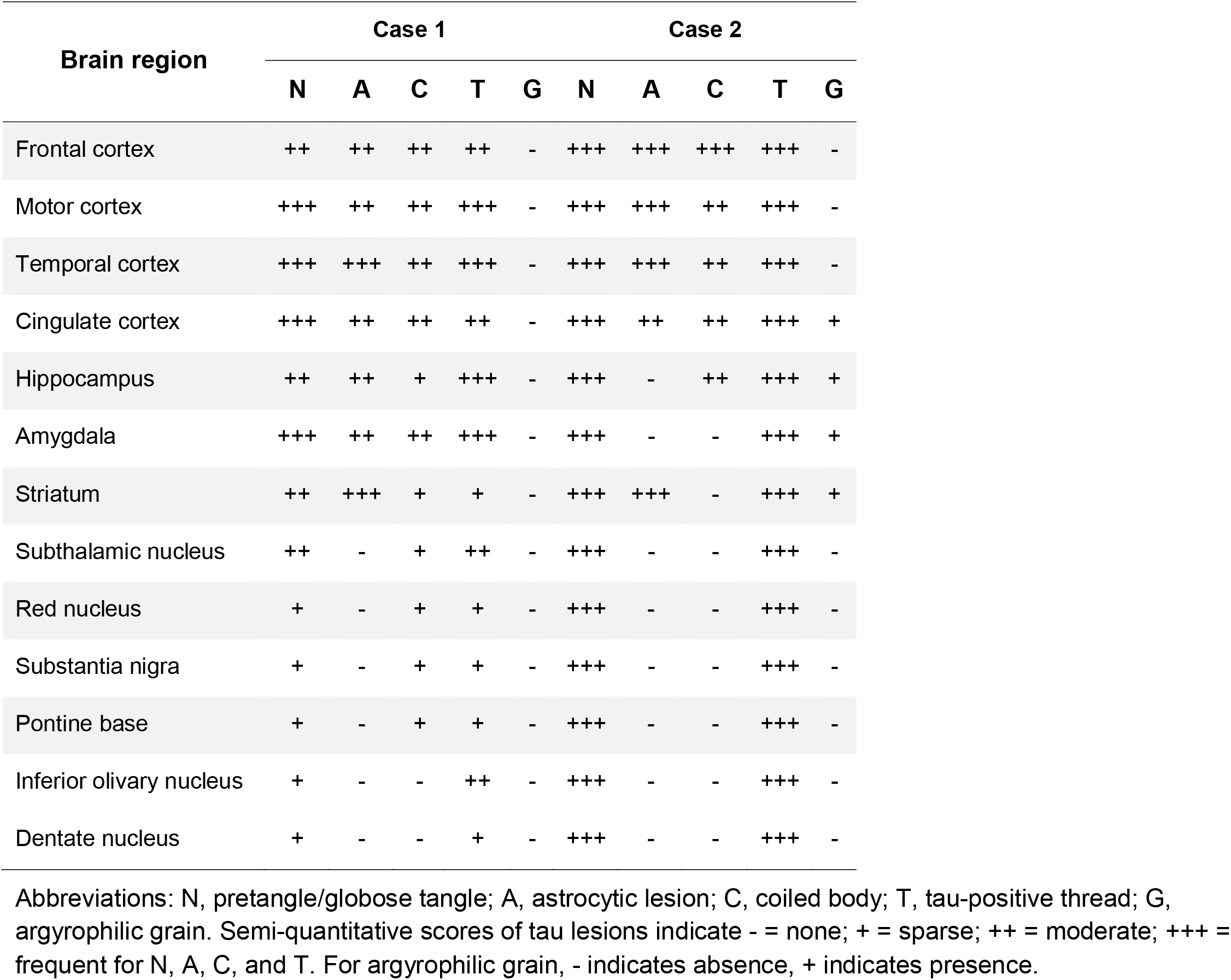
Topographical distribution of tau pathology.

#### Case 1

The fixed left hemibrain weighed 460 grams. The macroscopic findings revealed minimal cortical atrophy in frontal and temporal lobes, but there was thinning of the corpus callosum and dilation of the lateral ventricles. Subcortical regions, brainstem, and cerebellum were unremarkable except for decreased pigmentation in substantia nigra and locus coeruleus (**Figure 2A–D**).

**Figure 2:**
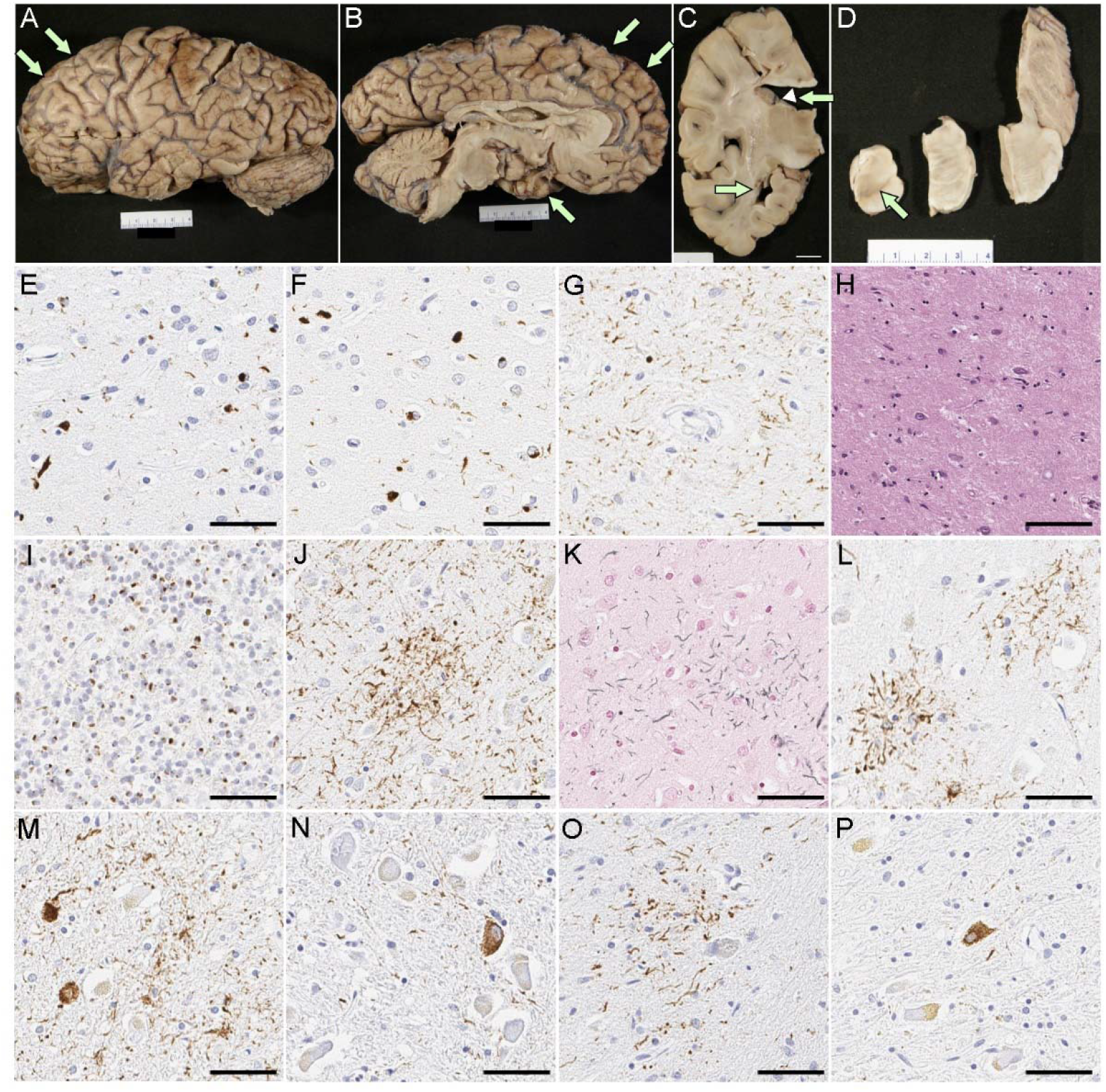
Pathologic findings of Case 1. (A-D) Macroscopic finding reveals minimal cortical atrophy in the frontal and temporal lobes (arrows in A and B), thinning of the corpus callosum (arrowhead in C), and marked enlargement of the frontal horn of the lateral ventricle (arrows in C). The substantia nigra shows loss of pigment (arrow in D). Immunohistochemistry for pTDP-43 (E-G), C9RANT (I), and CP13 (J, L-P), as well as H&E (H) and Gallyas staining (K). There are neuronal cytoplasmic inclusions, dystrophic neurites, and neuronal intranuclear inclusions in the midfrontal cortex (E) and inferior temporal cortex (F), consistent with FTLD-TDP type A. (G) CA1 sector of the hippocampus has neuronal cytoplasmic inclusions and numerous fine neurites, accompanied by neuronal loss (H), consistent with hippocampal sclerosis. (I) Numerous inclusions are C9rant-positive in the cerebellar granular layer. Astrocytic plaques are present in the motor cortex (J, K) and superior frontal cortex (L). A mild degree of pretangles and threads are present in the subthalamic nucleus (M), pontine base (N), inferior olivary nucleus (O), and cerebellar dentate nucleus (P). Scale bars: 1 cm in C, 50 μm in E-P.

On H&E-stained sections, the neocortex had mild spongiform change and gliosis, as well as mild thinning of the cortical ribbon, most marked in the frontal and temporal lobes. Only a few balloon neurons were detected in the cingulate gyrus. Immunohistochemistry for phospho-TDP-43 showed neuritic processes and neuronal cytoplasmic inclusions (NCI) in the neocortex, most marked in superficial cortical layers (**Figure 2E, F**), consistent with FTLD-TDP type A [31]. A few neuronal intranuclear inclusions were also detected. TDP-43-positive fine neurites were present in the CA1 sector of the hippocampus, along with marked neuronal loss, consistent with hippocampal sclerosis (**Figure 2G, H**). The caudate nucleus and the nucleus accumbens had atrophy and gliosis, while the putamen and globus pallidus were less affected. Sparse NCI and a dystrophic neurites were detected. The thalamus had mild atrophy and gliosis in anterior and dorsomedial nuclei. The ventral and lateral thalamus and the subthalamic nucleus were unremarkable. The substantia nigra had marked neuronal loss and gliosis with extraneuronal neuromelanin, marked in the ventrolateral cell group. Moderate numbers of NCI, including skein-like inclusions, were present. The cerebral peduncle had atrophy and myelinated fiber loss in the frontobulbar tract, but not the corticospinal tract. The raphe nuclei had a mild neuronal loss, but the locus coeruleus was well populated. The medullary pyramid and hypoglossal nucleus were unremarkable. The inferior olivary nucleus had a mild neuronal loss, but more marked gliosis and moderate NCI. Immunohistochemistry for C9RANT [32] revealed numerous neuronal inclusions in the cerebellar granular cell layers (**Figure 2I**), confirming *C9ORF72* mutation.

Phospho-tau immunohistochemistry and Gallyas staining revealed astrocytic plaques in the superior frontal gyrus and premotor cortex (**Figure 2J–L**). Moderate threads and coiled bodies were detected in adjacent white matter. Sparse tau pathology, mainly pretangles and threads, was also observed in the subthalamic nucleus, pontine base, inferior olivary nucleus, and cerebellar dentate nucleus (**Figure 2M–P**).

#### Case 2

The fixed left hemibrain weighed 540 grams. Macroscopic evaluation of the fixed brain revealed severe cortical atrophy over the dorsolateral and medial frontal lobe, including the frontal pole and the orbital frontal lobe (**Figure 3A, B**). The medial temporal lobe had moderate atrophy and the parietal lobe had mild atrophy in the superior lobule. The anterior corpus callosum was markedly thinned (**Figure 3C**). The hippocampal formation and amygdala were both atrophic, especially in the subiculum. Basal ganglia showed severe atrophy of the caudate nucleus and attenuation of the anterior limb of the internal capsule (**Figure 3C**). The globus pallidus was markedly atrophic and had brown discoloration. The anterior thalamus was atrophic. The substantia nigra had marked loss of pigment (**Figure 3D**).

**Figure 3:**
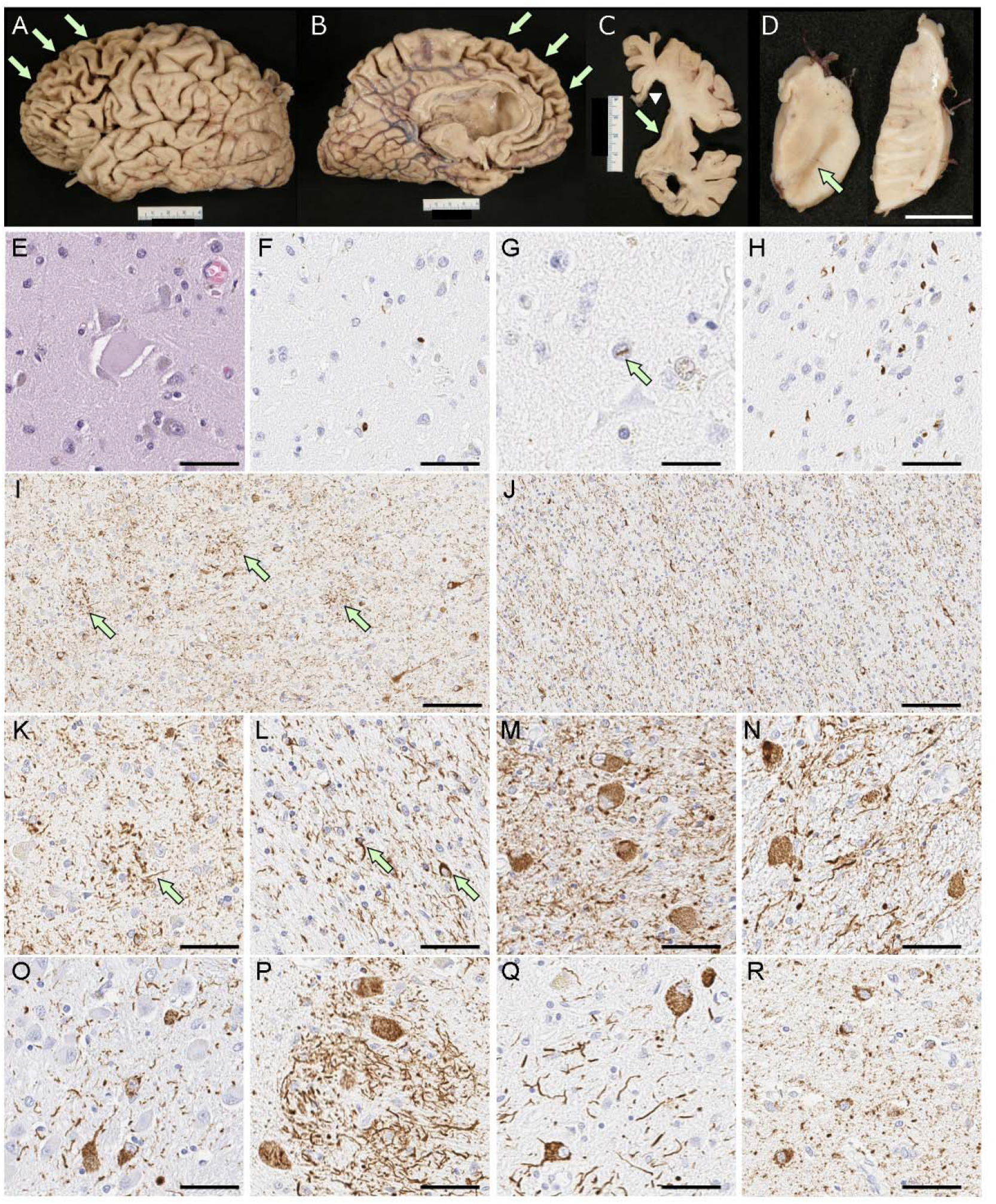
Pathologic findings of Case 2. (A-D) Macroscopic finding reveals severe cortical atrophy over the frontal lobe (arrows in A and B), thinning of the corpus callosum (arrowhead in C), and massive enlargement of the lateral ventricles, especially the frontal horn (C). Basal ganglia show severe atrophy of the caudate nucleus with a concave ventricular surface (arrow in C) and attenuation of the anterior limb of the internal capsule. The substantia nigra shows marked loss of pigment (arrow in D). Balloon neurons are observed in the superior frontal gyrus on H&E-stained sections (E). Immunohistochemistry for pTDP-43 shows neuronal cytoplasmic inclusions (F), dystrophic neurites, and neuronal intranuclear inclusions (G) in the superficial layer of the superior frontal cortex, consistent with FTLD-TDP type A. (H) The amygdala has neuronal cytoplasmic inclusions and dystrophic neurites. Immunohistochemistry for CP13 shows astrocytic plaques and numerous threads in the gray matter of the superior frontal gyrus (I, K). Numerous threads and coiled bodies are observed in the adjacent white matter (J, L). Abundant pretangles with threads are present in the subthalamic nucleus (M), red nucleus (N), pontine base (O), inferior olivary nucleus (P), and dentate nucleus (Q). Argyrophilic grains and pretangles are observed in the cingulate gyrus (R). Scale bars: 1 cm in D; 50 μm in E, G, H, and L-S; 25 μm in F; 100 μm in I and J.

On microscopic evaluation, the neocortex had thinning of the cortical ribbon with spongiform change and extensive gliosis, most marked in the frontal cortices, but sparing of the primary cortices and the occipital lobe. The gliosis was striking at the gray-white junction. The cingulate and frontal cortices had ballooned neurons (**Figure 3E**) on H&E-stained sections. Immunohistochemistry for phospho-TDP-43 revealed neuritic processes, NCI (**Figure 3F**), and a few intranuclear inclusions (**Figure 3G**) in the neocortex, most marked in the frontal cortex, amygdala (**Figure 3H**), dentate fascia, pyramidal layer of the hippocampus, substantia nigra, red nucleus, and inferior olivary nucleus. H&E staining showed severe neuronal loss and gliosis in CA1 and the subiculum. These findings were consistent with FTLD-TDP type A with hippocampal sclerosis [31]. There was also extensive atrophy of the basal ganglia and diffuse gliosis in the caudate nucleus and nucleus accumbens, while the globus pallidus was least affected. The anterior limb of the internal capsule had marked attenuation with myelinated fiber loss and gliosis. The thalamus, subthalamic nucleus, and adjacent hypothalamus had marked gliosis. The mammillary body was atrophic and had many pyknotic neurons consistent with transsynaptic degeneration. The posterior limb of the internal capsule was pale and gliotic. The substantia nigra had marked neuronal loss with extraneuronal neuromelanin and gliosis. Sparse TDP-43-positive NCI and dystrophic neurites were present. The cerebral peduncle had atrophy and gliosis of the frontobulbar tract. The raphe nuclei, locus coeruleus, reticular formation, medullary pyramid, and hypoglossal nucleus were histologically unremarkable. The inferior olivary nucleus had severe gliosis with many TDP-43-positive NCI.

Immunohistochemistry for phospho-tau revealed tau-positive threads and astrocytic plaques in the neocortex (**Figure 3I, K**), as well as numerous threads and fewer coiled bodies in cerebral white matter (**Figure 3J, L**) consistent with CBD. Abundant pretangles and threads were present in the subthalamic nucleus, red nucleus, pontine nuclei, inferior olivary nucleus, and dentate nucleus (**Figure 3M–Q**), but the neuronal populations in these regions were relatively well preserved. Argyrophilic grains and pretangles were observed in the amygdala, hippocampus, ventral striatum, and cingulate gyrus (**Figure 3R**), consistent with AGD.

## Discussion

In this series of 201 autopsy cases with TDP-43 proteinopathies, many patients had concurrent tau pathologies. As expected, ARTAG (42%) and PART (36%) were the most frequent tau pathologies, followed by AGD (22%). In addition, we found two patients with CBD, which is a 4-repeat tauopathy form of FTLD-tau. TDP-43 and tau are the most common molecular subtypes of FTLD, but coexistence of FTLD-TDP and FTLD-tau is uncommon [18]. These cases raise the issue of what should be considered the “primary” pathologic process, and which may be considered “secondary” or “co-primary” process.

Case 1 was clinically diagnosed with familial AD based upon his cognitive impairment, initially characterized by amnestic type dementia, as well as dementia in multiple family members. Neuropathologic assessment revealed mild Alzheimer’s-type pathology (Braak NFT stage III and Thal amyloid phase 3) insufficient to account for dementia. Although the cortical atrophy of the frontal and temporal lobes was relatively mild, immunohistochemistry for TDP-43 and presence of C9RANT inclusions were diagnostic of FTLD-TDP [32]. A hexanucleotide repeat expansion in the *C9ORF72* gene was confirmed with repeat-primed polymerase chain reaction assay [30]. The presence of a pathogenic mutation in a gene for FTLD makes a strong case for the primary diagnosis in the case to be FTLD-TDP. Of note, amnestic Alzheimer’s dementia is a common clinical diagnosis of genetically-confirmed FTLD-TDP in elderly individuals [33], and in an autopsy series from the State of Florida brain bank, late onset patients with *C9ORF72* mutations often present with Alzheimer type dementia or Lewy body dementia [34].

Tau pathology in the cases was consistent with CBD based upon morphology and neuroanatomical distribution; however, tau pathology was mild and subcortical nuclei vulnerable to neuronal loss in CBD, globus pallidus and substantia nigra, were well preserved. Moreover, the severity of tau pathology was mild in subcortical regions. These findings are similar to those reported as “preclinical” CBD [35]. In the current situation the term “preclinical” is not appropriate since the patient presented with cognitive impairment, behavioral changes, and parkinsonism. Taken together, genetically confirmed FTLD-TDP with “incidental” CBD seems to be the best neuropathologic diagnosis.

Interestingly, this patient is similar to a patient in the study of Kim and coworkers. Their patient had FTLD-TDP type A and unclassifiable FTLD-tau with *C9ORF72* mutation (Case 1) [18]. Given that *C9ORF72* mutation is the most common risk factor for familial amyotrophic lateral sclerosis and FTLD, its strong association with TDP-43 pathology has been established. In contrast, it is still unknown whether *C9ORF72* mutation is also associated with FTLD-tau. Bieniek and colleagues investigated Alzheimer’s-type tau pathology in the temporal cortex and hippocampus in patients with FTLD who carried *C9ORF72* mutation (c9FTLD) [36], and found that tau pathology burden was not different between c9FTLD and sporadic FTLD. Snowden and colleagues screened for *C9ORF72* mutations in 398 patients with clinical presentations of FTD and found one patient who had CBD pathology without TDP-43 pathology [37]. The absence of TDP-43 pathology raised the question of the significance of *C9ORF72* hexanucleotide expansion in this patient. King and colleagues reported a patient with *C9ORF72* mutation who had Pick’s disease and TDP-43 pathology, with TDP-43 co-localizing with Pick bodies [38]. This patient also had a *MAPT* variant A239T, which has not been associated with tau pathology. Taken together, the association between *C9ORF72* mutation and tau pathology remains uncertain and warrants further investigation.

Our second case had marked atrophy in frontal and temporal lobes, as well as severe atrophy in the orbital gyrus and caudate nucleus, and hippocampal sclerosis consistent with FTLD-TDP. In contrast, marked pigment loss in the substantia nigra and discoloration of the globus pallidus were suggestive of CBD. TDP-43 immunohistochemistry revealed moderate NCI and DN and sparse neuronal intranuclear inclusions in the neocortex, consistent with FTLD-TDP type A. There was also severe tau pathology in the grey and white matter of the neocortex, deep gray nuclei, brainstem, and cerebellum, consistent with CBD. Given that both FTLD-TDP and CBD can present as behavioral variant frontotemporal dementia, it is difficult to determine which pathology is “primary” for this patient. One can argue the that CBD is the “primary” diagnosis in this patient, and that TDP-43 pathology can be explained by limbic-predominant age-related TDP-43 encephalopathy neuropathological change (LATE-NC) [39]. LATE-NC is characterized by TDP-43 proteinopathy in the medial temporal lobe in elderly with or without hippocampal sclerosis [39]. The amygdala and hippocampus are the most vulnerable regions, but the midfrontal gyrus is affected in stage 3; therefore, it is challenging to differentiate FTLD-TDP and advanced LATE-NC. Case 2 had significant frontal lobe atrophy, which supports the diagnosis of FTLD-TDP; however, frontal lobe atrophy is also common in CBD. A diagnosis of CBD with LATE-NC is reasonable. The advanced age of the patient (84 years) and the presence of hippocampal sclerosis may also support the diagnosis of LATE-NC. Nevertheless, our final neuropathological diagnosis is FTLD-TDP with “incidental” CBD because atrophy of the globus pallidus and subthalamic nucleus were relatively mild compared to that of the orbital gyrus and caudate nucleus. We judge the CBD to be “incidental” pathology in the context of FTLD-TDP.

The present study needs to be taken account of the fact that TDP-43 pathology can be found in a subset of CBD cases. In our previous study, we identified astrocytic plaque-like TDP-43 lesions in the motor cortex and superior frontal gyrus in CBD patients with TDP-43 pathology. Therefore, we assumed that TDP-43 pathology was “secondary” to CBD pathology [17]. In contrast, the two patients in the present study did not have astrocytic plaques with TDP-43 immunoreactive processes. This finding supports the conclusion that TDP-43 pathology occurs independent of tau pathology in CBD, not “secondary” to tau pathology

Recently, Llibre-Guerra and colleagues reported a novel neuroglial tauopathy associated with transmembrane protein 106B (*TMEM106B*) rs1990622 A/A genotype in FTLD/ALS-TDP [40]. TMEM106B rs1990622 is a common variant in *TMEM106B* that increases the risk of FTLD-TDP, especially those caused by *GRN* mutations [41, 42]. This tauopathy was characterized by limbic-predominant distribution of grains, granular neuronal cytoplasmic inclusions, as well as diffuse granular immunopositivity in astrocytic processes and threads. Although these features resemble AGD or ARTAG, the distribution and overall morphological features were not consistent with these diagnoses. In the present study of tau in FTLD-TDP, the most common tau pathologies were ARTAG and AGD. We did not identify tau lesions similar to those reported as TMEM106B-associated neuroglial tauopathy.

A limitation of this study is that the clinical information of two patients with FTLD-TDP and CBD was limited due to the retrospective nature of the autopsy cohort. Systematic, longitudinal data with neurological examinations and neuropsychological testing were not available. Motor symptoms suggestive of corticobasal syndrome may have been overlooked or not well documented. Another limitation is that we did not assess clinical features in patients with PART, ARTAG, or AGD because the primary focus was FTLD-tau, rather than these “age-related” tau pathologies.

In conclusion, by screening for tau pathology in TDP-43 proteinopathies, we identified two patients with mixed FTLD-TDP and CBD that are likely independent (“co-primary”) disease processes. Many of elderly individuals with neurodegenerative disorders have multiple coexisting pathologies [8, 10]; therefore, both TDP-43 and tau should be part of the screening process for the neuropathologic assessment of FTLD. We are not able to determine the relative contribution of each pathology to the clinical presentations in these patient. This is not too dissimilar to the issue of assigning relative contributions of mixed pathology in other settings, for example, cases with both Alzheimer’s disease and diffuse Lewy body disease. Implementation of molecular imaging modalities for each protein may help determine the timing and relative contributions of the proteins to clinical presentations.

## Acknowledgements

We would like to thank the patients and their families who donated brains to help further the scientific understanding of neurodegeneration. The authors would also like to acknowledge Virginia Phillips, Jo A. Landino Garcia, and Ariston L. Librero (Mayo Clinic, Jacksonville) for histologic support, and Monica Castanedes-Casey (Mayo Clinic, Jacksonville) for immunohistochemistry support. This work is supported by CurePSP, the Rainwater Charitable Trust, and the Jaye F. and Betty F. Dyer Foundation Fellowship in progressive supranuclear palsy research, as well as NINDS Tau Center without Walls (U54-NS100693).

## Ethical statement

Brain autopsies were obtained after consent of the legal next-of-kin or individuals with legal authority to grant autopsy permission. De-identified studies of autopsy samples are considered exempt from human subject research by the Mayo Clinic Institutional Review Board.

## Author Contributions

**Shunsuke Koga**: Study concept and design; acquisition, analysis, and interpretation of data; execution of the statistical analysis; drafting of the manuscript. **Xiaolai Zhou**: Study concept and design; acquisition, analysis, and interpretation of data; review and critique. **Aya Murakami**: Acquisition, analysis, and interpretation of data; review and critique. **Cristhoper Fernandez De Castro**: Acquisition of data; review and critique. **Matthew C. Baker**: Acquisition, analysis, and interpretation of data; review and critique. **Rosa Rademakers**: Review and critique. **Dennis W. Dickson**: Study concept and design; acquisition and interpretation of data; review and critique.

## Data Availability Statement

The data that support the findings of this study are available from the corresponding author upon reasonable request.

## Supplementary Data: Clinical history of two patients with FTLD-TDP and CBD

Case 1: This patient was an 89-year-old, right-handed Hispanic man with an 8-year history of memory impairment and a 3-year history of disorientation, socially inappropriate behavior, and apathy. His family history was notable for late-onset (70s to 80s) dementia in his father and two siblings who died in the 80s to 90s. Neurological examination revealed saccadic eye movements, action tremor in both upper extremities, mild bradykinesia in both lower extremities, and a slow unsteady gait. He scored 9/30 on the Mini-Mental State Examination. He was diagnosed with familial Alzheimer’s disease. His dementia and gait difficulty gradually worsened. He developed frequent falls in the last 6 months of his life.

Case 2: This 84-year-old Caucasian man had a six-year history of memory loss and behavior change, characterized by agitation, mutism and violent outbursts, treated with rivastigmine, memantine, and risperidone. He had no family history of neurological disorders. On neurological examination at 78 years of age, he scored 15/30 on the Mini-Mental State Examination. An MRI of the brain revealed mild cortical atrophy, and FDG-PET revealed decreased glucose metabolism in the bilateral frontal and temporal lobes. He was diagnosed with frontotemporal dementia with a differential diagnosis of mixed dementia (i.e., Alzheimer’s-type, vascular, and alcohol-related dementia).

